## Supplemental information for "Towards coevolution-aware ancestral sequence reconstruction"

### Supplementary Material for
### "Towards coevolution-aware ancestral sequence reconstruction"

Alya Zeinaty,[1,2] Leonardo di Bari,[1,3] Saverio Rossi,[2]
Pierre Barrat-Charlaix,[1] Francesco Zamponi,[2,*] and Martin Weigt[1,4,*]

[1] *Sorbonne Université, CNRS, Computational, Quantitative and Synthetic Biology, 75005 Paris, France*
[2] *Dipartimento di Fisica, Sapienza Università di Roma, Piazzale Aldo Moro 5, 00185 Rome, Italy*
[3] *DISAT, Politecnico di Torino, Corso Duca degli Abruzzi, 24, I-10129, Torino, Italy*
[4] *Institut Universitaire de France (IUF)*

(Dated: June 8, 2026)

#### I. SEQUENCE DATA PROCESSING

Our procedure takes as input an MSA of extant protein sequences $\mathcal{D}_{\text{extant}}$, but is also applicable, with minor modifications, to RNA or DNA alignments. For clarity purposes, the rest of the text will however focus on protein alignments. In our analysis, sequence alignments were obtained by scanning UniProt using *hmmer* [1], starting from a set of sequences, or *seed*, representative of the protein family of interest. Sequences containing more than a fixed fraction of gap characters (20% in all analyses presented here) are then discarded, to improve quality and reduce noise introduced by poorly aligned or fragmentary sequences. Following this initial filtering, duplicate sequences are removed using a gap-aware removal procedure. Two sequences are considered equivalent if they are identical at all positions in which both have a non-gap character. For each group of equivalent sequences, a single representative is retained, chosen as the sequence with the minimal number of gaps. This strategy removes exact duplicates while preferentially preserving sequences with maximal information content, without introducing an explicit sequence identity threshold or clustering radius. The resulting alignment $\mathcal{D}_{\text{extant}}$ constitutes the final sequence dataset used for phylogenetic inference.

The alignment provided as input for ancestral sequence inference and the learning of Direct Coupling Analysis parameters must satisfy specific properties to ensure the mathematical validity of the resulting model. First, it is essential that all sequences are aligned to a uniform length $L$. Furthermore, the alignment must contain sufficient information to allow for the learning of $O((L \cdot q)^2)$ parameters for the DCA model, where $q$ is the alphabet size encompassing the 20 standard amino acids and the gap character, such that $q = |\{\text{amino acids}\}| + |\{\text{gap}\}| = 21$.

#### II. PHYLOGENETIC TREE INFERENCE

The filtered alignment obtained above is then used to infer the phylogenetic tree of the corresponding protein family. Several tools for phylogenetic inference have been developed over the past decade, including IQ-TREE [2], RAxML [3], and PhyML [4], which reconstruct trees by maximum likelihood using stochastic optimization procedures. More recently, Nesterenko *et al.* introduced PhyloFormer [5], a neural-network-based approach to phylogenetic inference. However, many of these methods are primarily designed for relatively shallow phylogenies involving a few hundred sequences, and become computationally expensive or inefficient when applied to large alignments with $M \approx 10^4$ sequences.

For this reason, we infer phylogenies using FastTree [6], which provides a favorable compromise between speed and accuracy for large protein families. FastTree first explores tree topologies using heuristic neighbor-joining, then refines them through a combination of nearest-neighbor interchanges and subtree prune–regraft moves to reduce total tree length. Final optimization is performed using maximum-likelihood rearrangements under a specified substitution model. In our analyses, we employ the Whelan–Goldman model for amino acid substitutions [7], include a pseudocount to improve numerical stability, and account for rate heterogeneity across sites by assigning sites to 20 discrete Gamma-distributed rate categories. To maximize numerical accuracy, FastTree is run in double precision with the options `-wag -pseudo -gamma`.

While this approach yields a robust tree at low computational cost, the resulting phylogeny typically contains a substantial number of branches with quasi-zero length (on the order of $10^{-8}$), reflecting unresolved relationships.

For clarity and to avoid redundancy, we apply a post-processing pipeline that collapses branches below a fixed length threshold ($10^{-6}$) by merging the corresponding child node with its parent. Internal nodes with a single child are also removed by promoting the child and summing branch lengths, thereby eliminating spurious linear chains while preserving total evolutionary distances (Supplementary Fig. 1). Since this procedure can affect leaf nodes, the alignment corresponding to the cleaned tree is stored separately as `family_name_collapsed_no_only_child.fasta`.

To verify that this cleaning step does not distort the global structure of sequence space, we compare principal component projections of the original and cleaned alignments, observing no substantial loss of information (panel F of supplementary figure 1). Finally, because FastTree assigns the root arbitrarily, the cleaned tree is rerooted at its midpoint using the `ete3` Python library via `t.get_midpoint_outgroup()` [8]. The resulting distribution of root-to-leaf distances is shown in supplementary figure 1. The final cleaned alignment and midpoint-rooted tree constitute the reference dataset used in all subsequent analyses.

#### III. SUPPLEMENTARY FIGURES

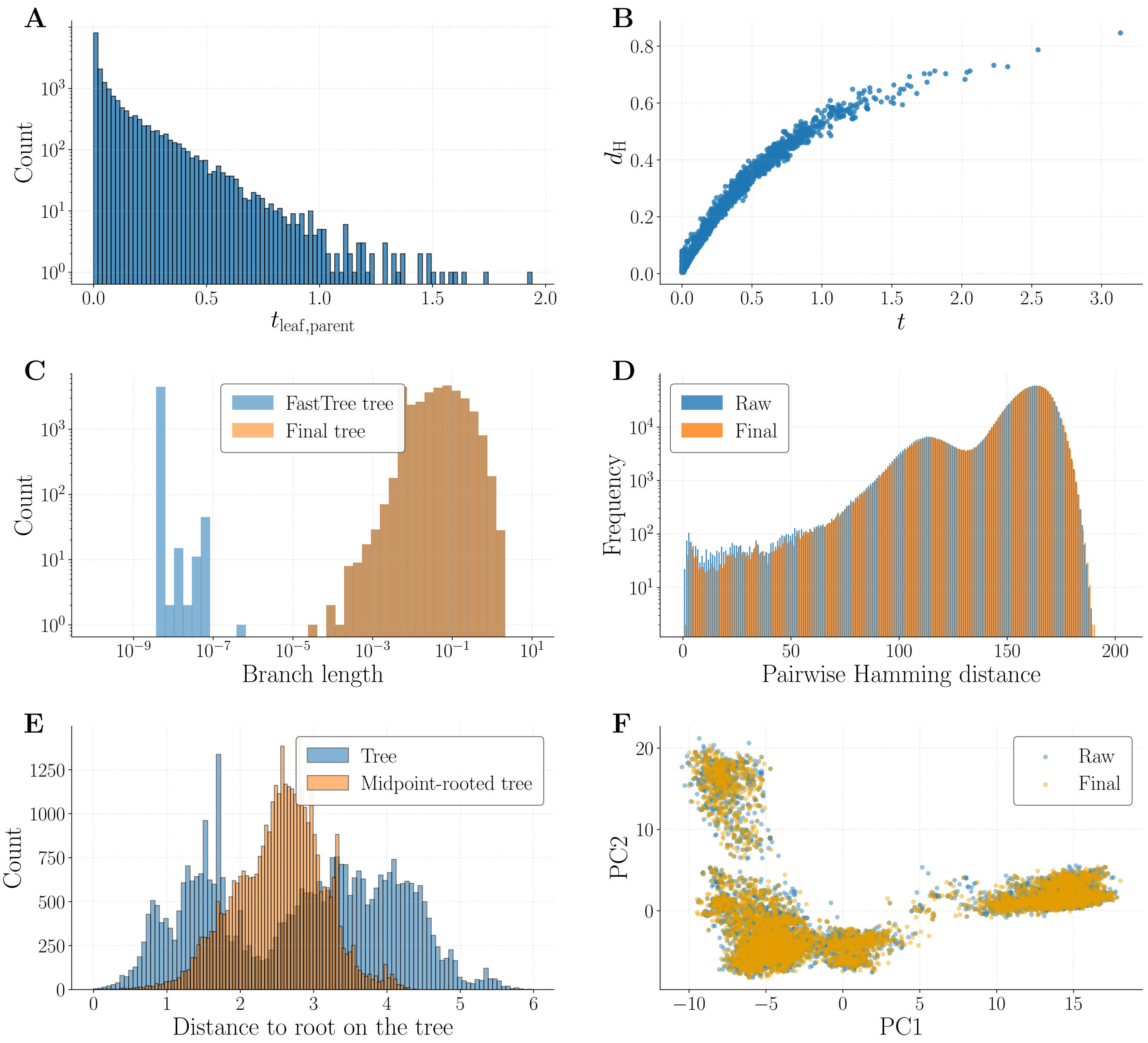


Supplementary Figure 1. **Tree and alignment polishing for the $\beta$-lactamase family** (A) $t_{\text{leaf,parent}}$ on the clean tree. (B) $d_{\text{H}}$ versus $t$ for any pair of leaves (tree: clean tree; data: cleaned alignment). (C) Branch-length distributions comparing the clean tree and the collapsed midpoint-rooted tree (tree-only analysis). (D) Pairwise Hamming distance distribution $d_{\text{H}}$ between the original raw alignment (no duplicates, gap percentage $< 20\%$) and the 'collapsed' alignment. (E) Distance to root on the clean and midpoint-rooted trees (data: cleaned alignment). (F) PCA of extant sequences projected from the raw and collapsed alignments.

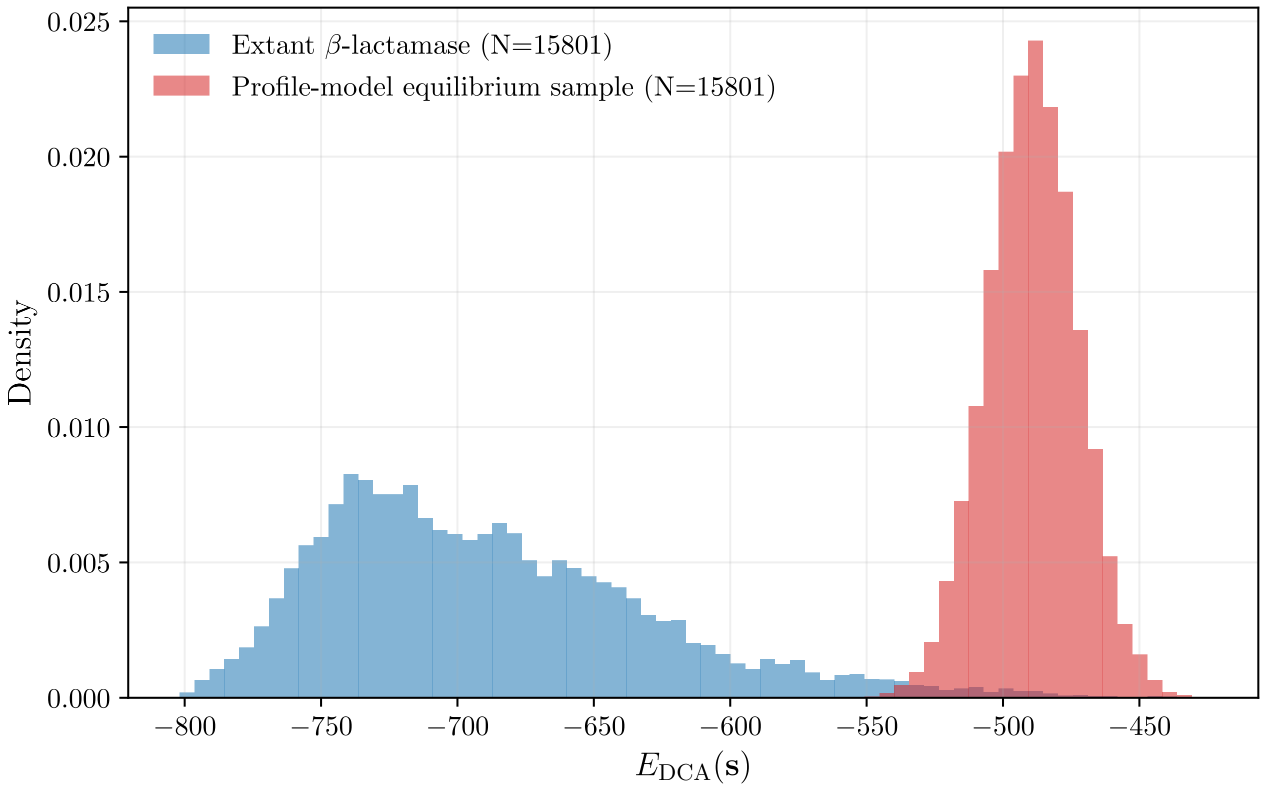


Supplementary Figure 2. **Comparison of DCA energies of the extant MSA and the profile-sampled MSA ($\beta$-lactamase)** Sequences sampled from the profile plotted against sequences from the extant MSA. 'Profile' means the probabilities taken from the single-site frequencies of the extant sequences. Energy distributions are evaluated by the DCA Potts model through the DCA energy $E_{\mathrm{DCA}}$. A low energy indicates 'good' sequences, while a high energy signifies highly atypical sequences, unlikely to be phenotypically functional according to the DCA model.

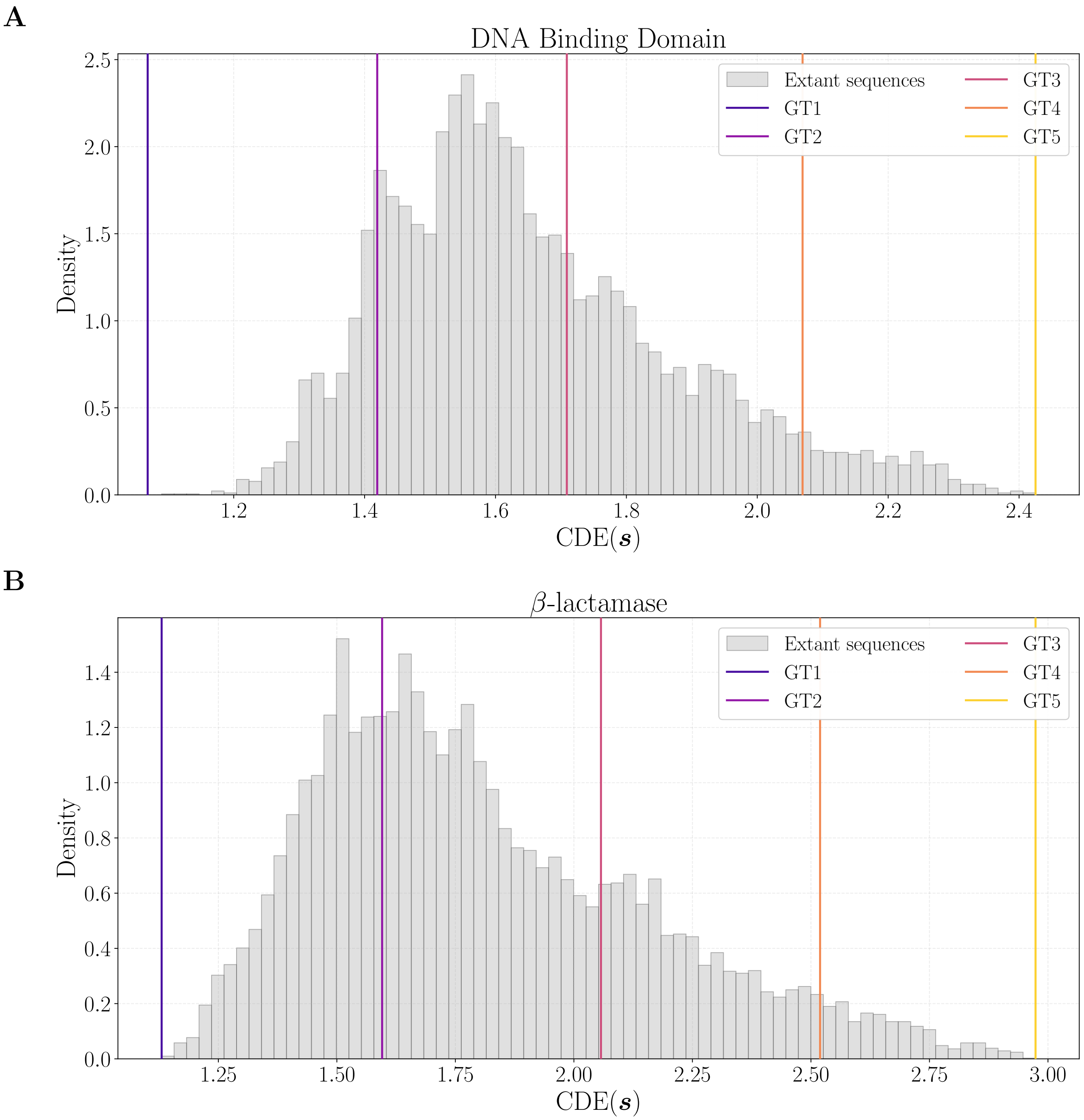


Supplementary Figure 3. **Context dependent entropy distribution of extant sequences** All extant sequences are plotted. On top, vertical lines for each of the sequences chosen as GT roots in the benchmarking procedure. (A) In the DNA Binding Domain (DBD) family. (B) In the $\beta$-lactamase family.

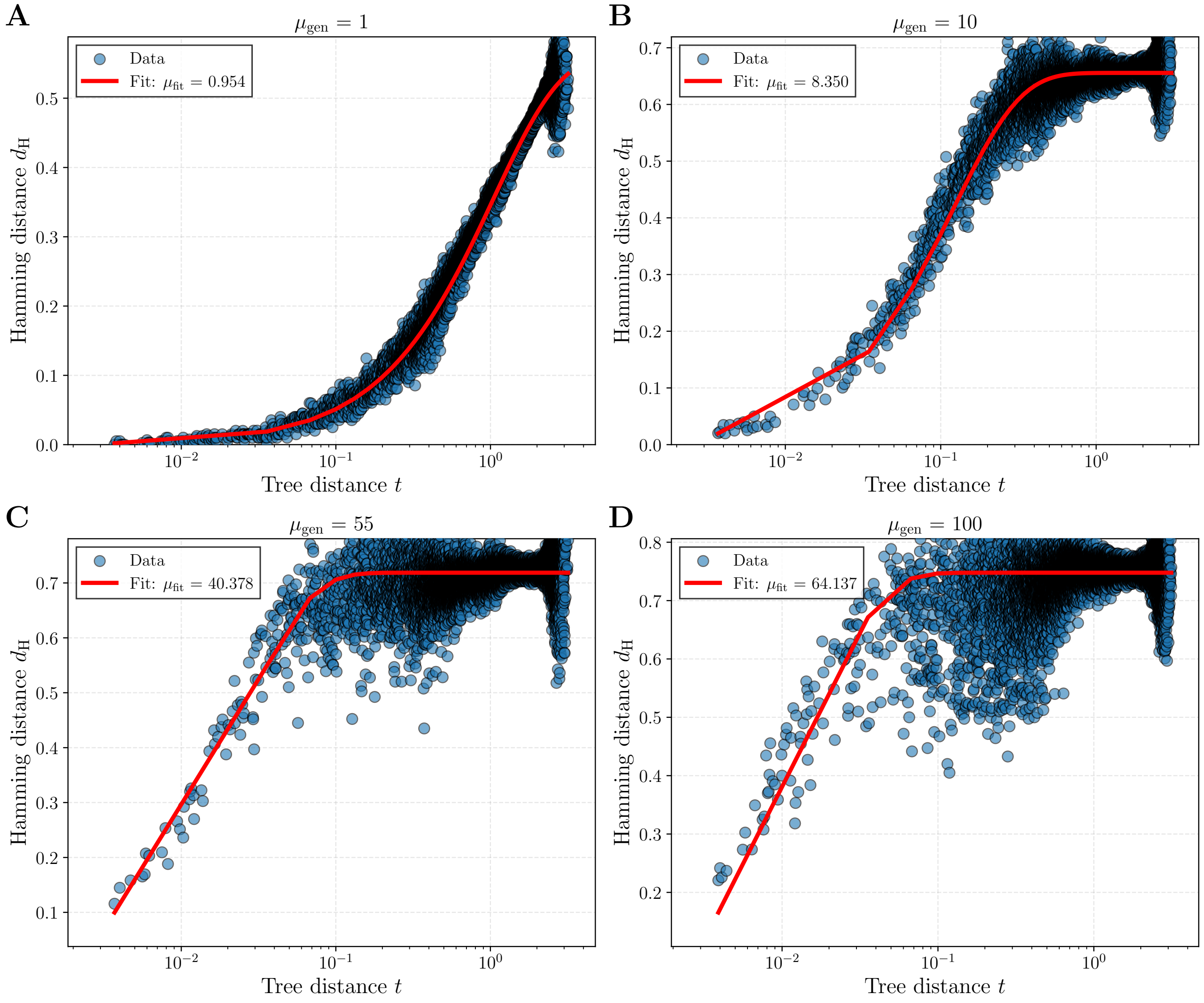


Supplementary Figure 4. **Curves of $\mu$ inference.** For each pair of sequences at the leaves, we plot the Hamming distance as a function of the time separating them on the tree. The best fit of the global mutation rate given the data $\boldsymbol{\mathcal{D}}$ at the leaves and the tree $\mathcal{T}$ is shown, inferred by fitting $\mathbb{E}[d_H(t)] = a(1 - e^{-\mu t})$. Here, $\mu_{\text{gen}}$ is the $\mu_{\text{gen}}$ of the DCA model, and $\mu_{\text{fit}}$ is the value of $\mu$ obtained after fitting. Evaluated on WT3 of the $\beta$-lactamase family.

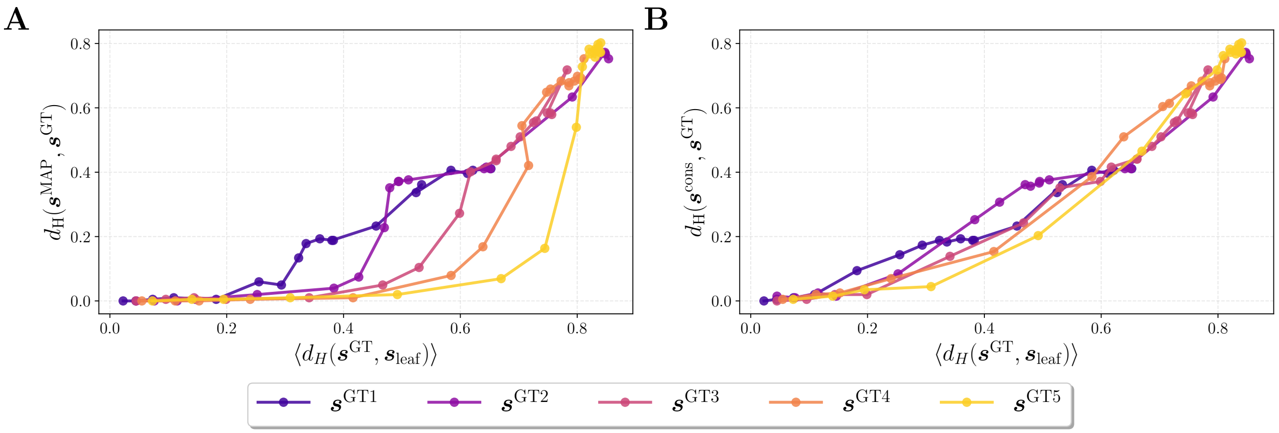


Supplementary Figure 5. **Another perspective on Figure 3 of the main text (on the $\beta$-lactamase family).** (A) Normalized Hamming distance between MAP and GT ancestors, as a function of average divergence between root and leaves. (B) Normalized Hamming distance between GT ancestor and consensus, as a function of average divergence between root and leaves.

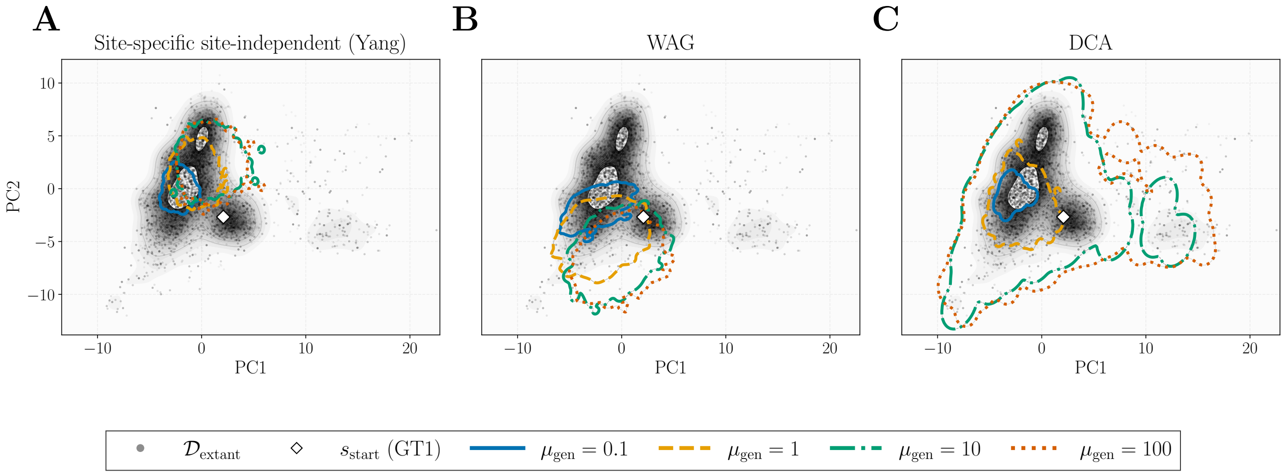


Supplementary Figure 6. **Exploration of the PCA space for 3 forward samplers, on the $\beta$-lactamase family.** (A) The Yang-defined site-specific and site-independent propagator. (B) The classical Whelan and Goldman transition model. (C) Our forward, coevolution-aware DCA-based simulator.

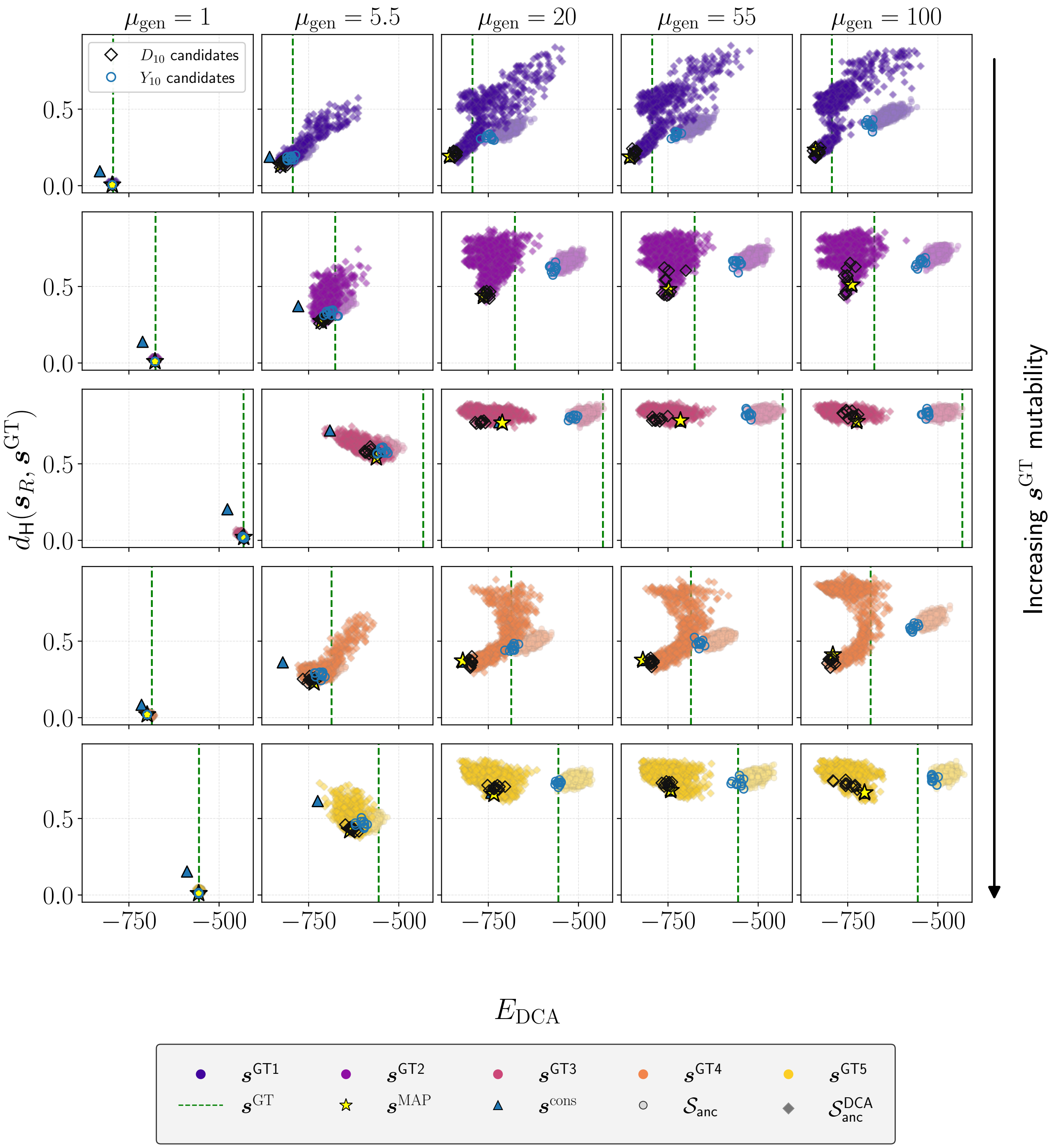


Supplementary Figure 7. $\mathcal{N} = 10$ **best candidates for each method.** On the $\beta$-lactamase family, plotted the $\mathcal{N} = 10$ best ancestral candidates according to the Yang site-independent likelihood $\mathbb{P}_R$, from both the site-independent ancestral set $\boldsymbol{\mathcal{S}}_{\text{anc}}$ ($Y_{10}$) and the 'reshuffled' coevolution-aware ancestral set $\boldsymbol{\mathcal{S}}_{\text{anc}}^{\text{DCA}}$ ($D_{10}$). Green dotted line represents the GT sequence $\boldsymbol{s}^{GT}$ (real root sequence), while the star is the MAP sequence $\boldsymbol{s}^{\text{MAP}}$ and the triangle is the consensus sequence of the leaves $\boldsymbol{s}^{\text{cons}}$. Plotted as a function of the DCA energy $E_{\text{DCA}}$ on the x-axis, and the normalized Hamming distance between candidate sequence and GT (y-axis).

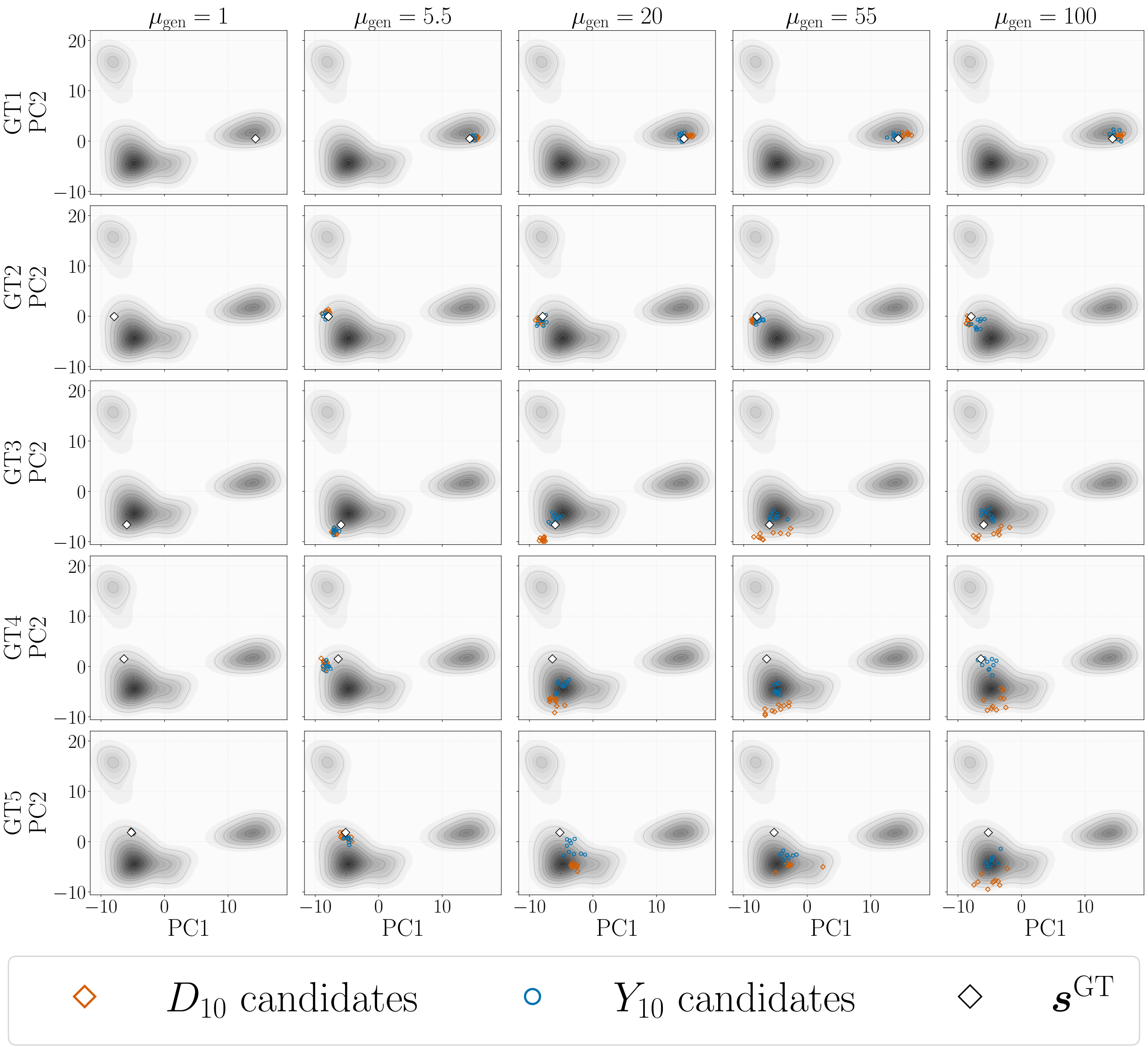


Supplementary Figure 8. $\mathcal{N} = 10$ **best candidates for each method, plotted on the PCA space of the extant sequences.** On the $\beta$-lactamase family, plotted the $\mathcal{N} = 10$ best ancestral candidates according to the Yang site-independent likelihood $\mathbb{P}_R$, from both the site-independent ancestral set $\boldsymbol{\mathcal{S}}_{\text{anc}}$ ($Y_{10}$) and the 'reshuffled' coevolution-aware ancestral set $\boldsymbol{\mathcal{S}}_{\text{anc}}^{\text{DCA}}$ ($D_{10}$). Projected on the Principal Components 1 and 2 of the extant alignment $\boldsymbol{\mathcal{D}}_{\text{extant}}$, represented by the grey density plot.

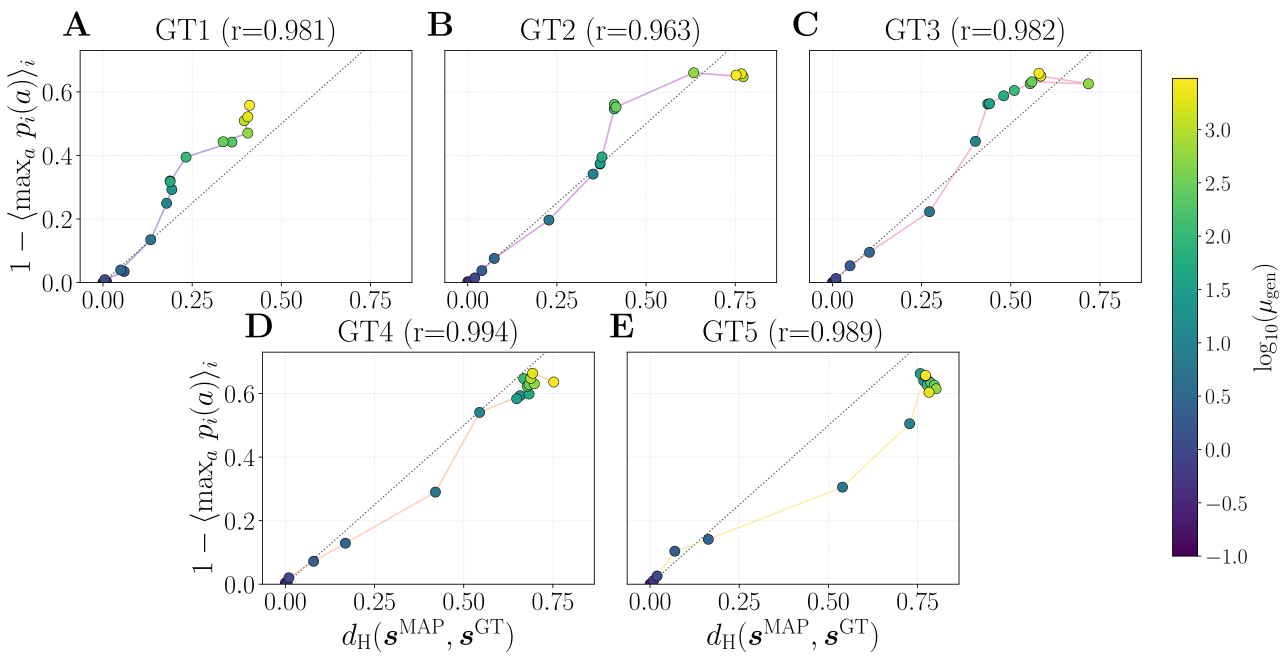


Supplementary Figure 9. **Correlation between Yang likelihood and Hamming distance to GT for the MAP sequence.** For each GT sequence of the $\beta$-lactamase family, plotted 1 - (average probability of the MAP sequence) against the Hamming distance to GT. The value r plotted as suptitle is the Pearson correlation between both metrics.

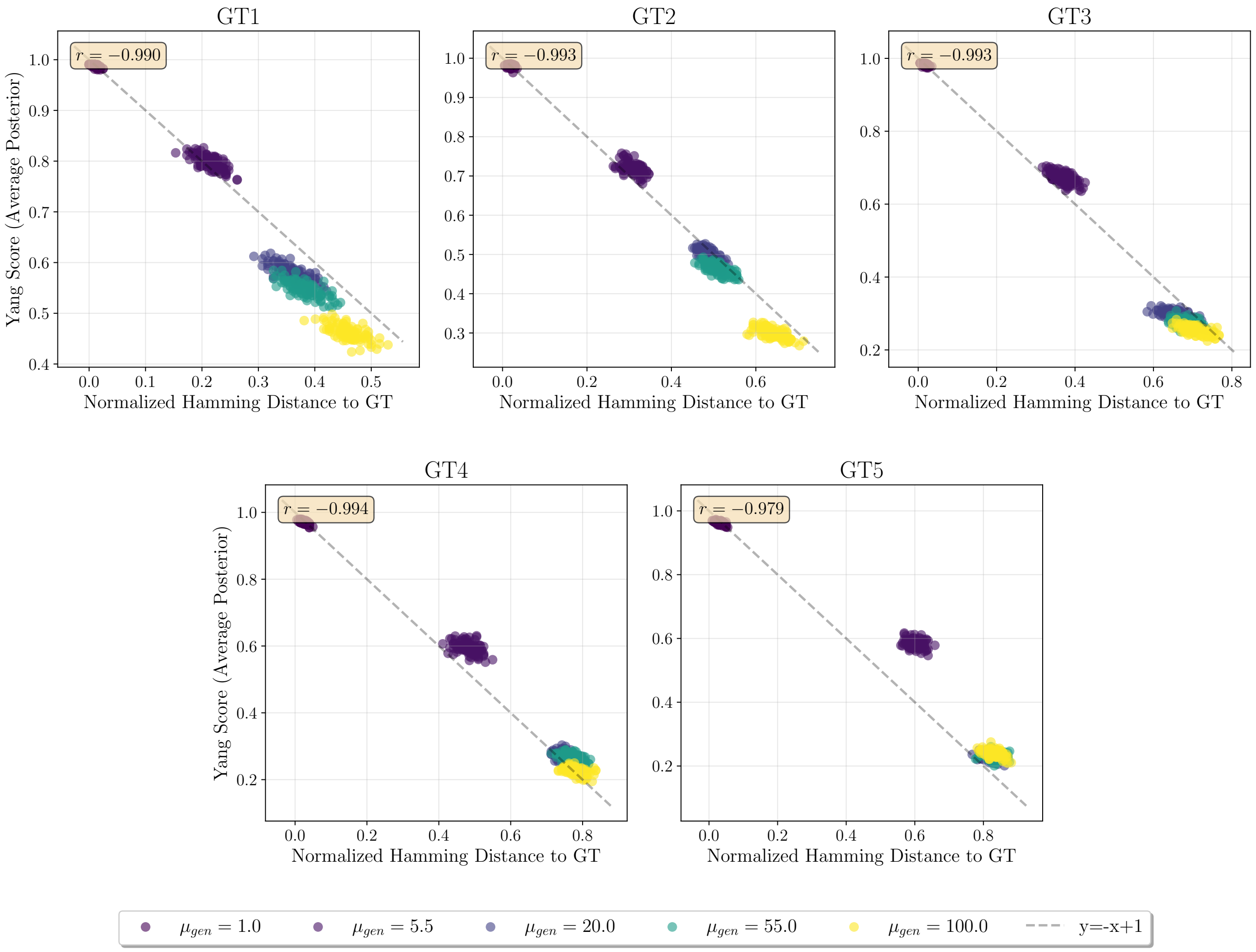


Supplementary Figure 10. **Correlation between Yang likelihood and Hamming distance to GT on site-independent ancestral samples.** For each GT sequence of the $\beta$-lactamase family, plotted $\mathbb{P}_R(\boldsymbol{s}_R)$ for sampled root sequences of $\boldsymbol{\mathcal{S}}_{\text{anc}}$, against the Hamming distance to GT. Plotted reconstructed sequences for different values of $\mu_{\text{gen}}$. The value r plotted as sup-title is the Pearson correlation between both metrics.

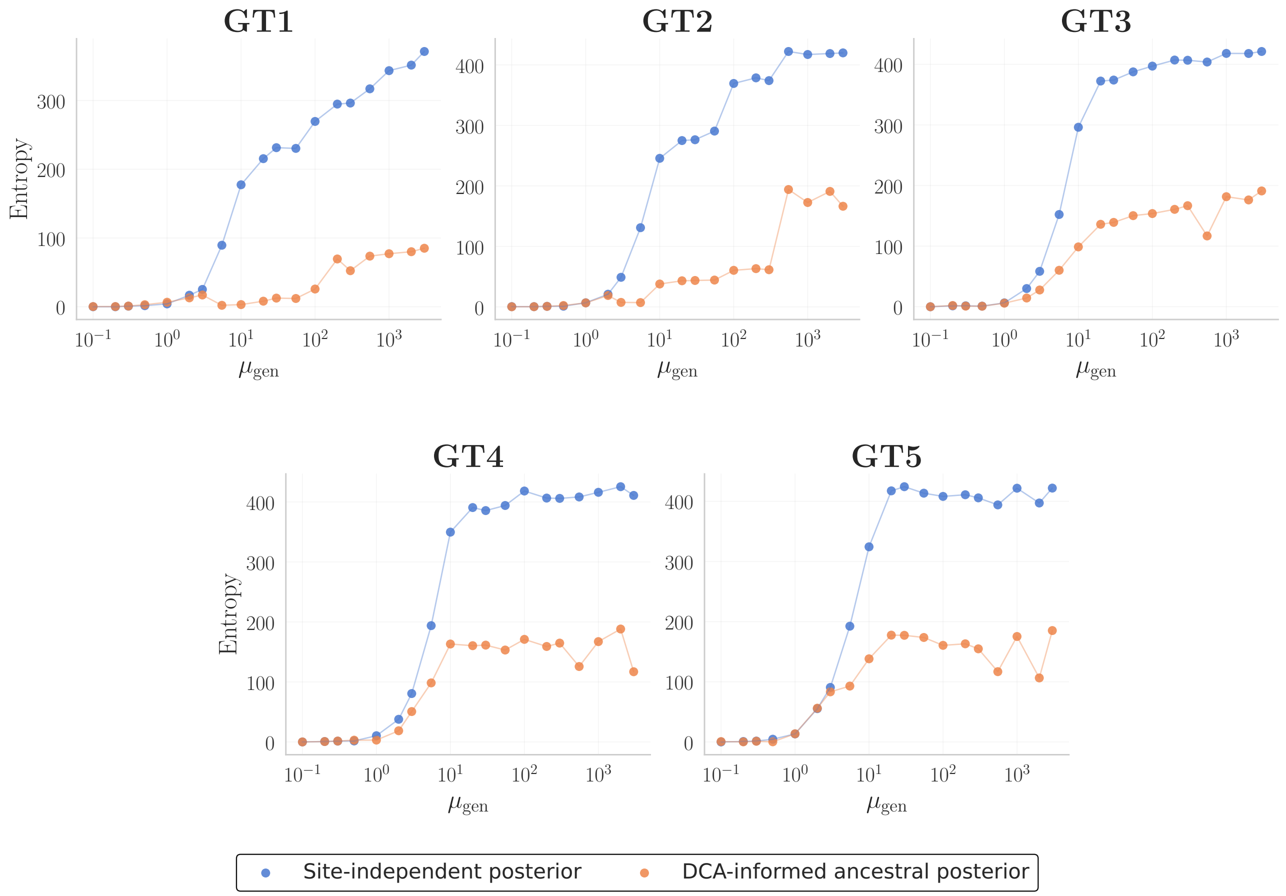


Supplementary Figure 11. **Measure of ancestral entropy on the $\beta$-lactamase family, with and without the coevolution couplings.** Site-independent entropy (entropy of the site-independent posterior $\mathbb{P}_R$, informed by the site-independent prior $\boldsymbol{\pi}$, i.e. the single-point frequencies taken from the leaves) was computed according to the standard entropy definition, and summed over sites. Entropy of the ancestral DCA model (site-independent conditional ancestral distribution, informed by a DCA prior) was computed approximately using Markov Chain Monte Carlo, with help from the adabmDCA [9] Git package. As expected, ancestral entropy is systematically reduced when couplings between sites are added as a prior to the ancestral distribution.

#### IV. RESULTS ON DBD

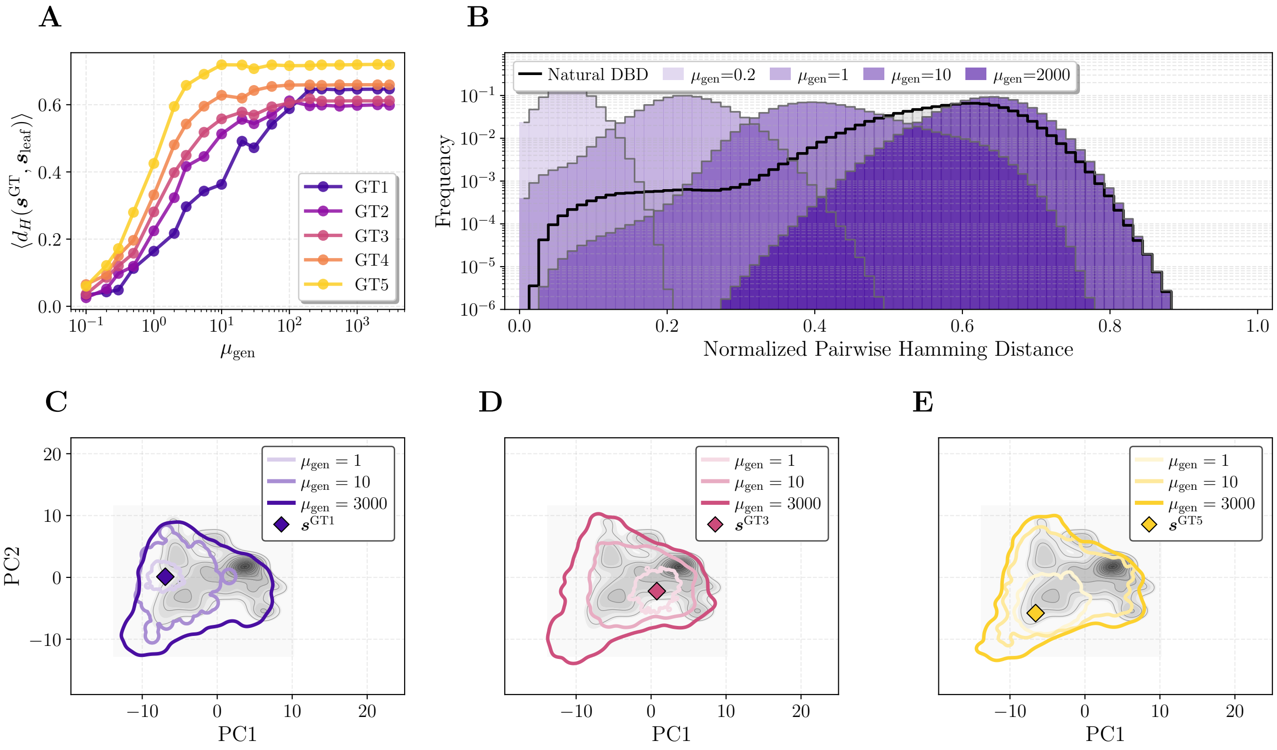


Supplementary Figure 12. **Effect of sampling time on forward generated sequences (DBD family).** Sequences were simulated with the DCA model along the phylogenetic tree of the DNA binding domain family, using increasing $\mu_{\rm gen}$ values (number of MCMC sweeps). (A) Average leaf–root Hamming distance divergence for simulated data as a function of $\mu_{\rm gen}$. (B) Distribution of pairwise Hamming distances $d_{\rm H}(\boldsymbol{s}^i, \boldsymbol{s}^j)$, with $\boldsymbol{s}^i, \boldsymbol{s}^j \in \boldsymbol{\mathcal{D}}$ among simulated leaves (shown here for $\boldsymbol{s}^{GT\,1}$). The gray histogram outlined in black represents the pairwise Hamming distance distribution of the extant sequences of the DBD family. (C,D,E) Principal Component Analysis of the sampled $\boldsymbol{\mathcal{D}}$ for $\boldsymbol{s}^{GT\,1}, \boldsymbol{s}^{GT\,3}, \boldsymbol{s}^{GT\,5}$ respectively. Extant sequences $\boldsymbol{\mathcal{D}}_{\rm extant}$ are plotted as the gray density distribution, on Principal Components 1 and 2. Coloured outlines show the exploration of PCA space for different evolutionary scales $\mu_{\rm gen}$.

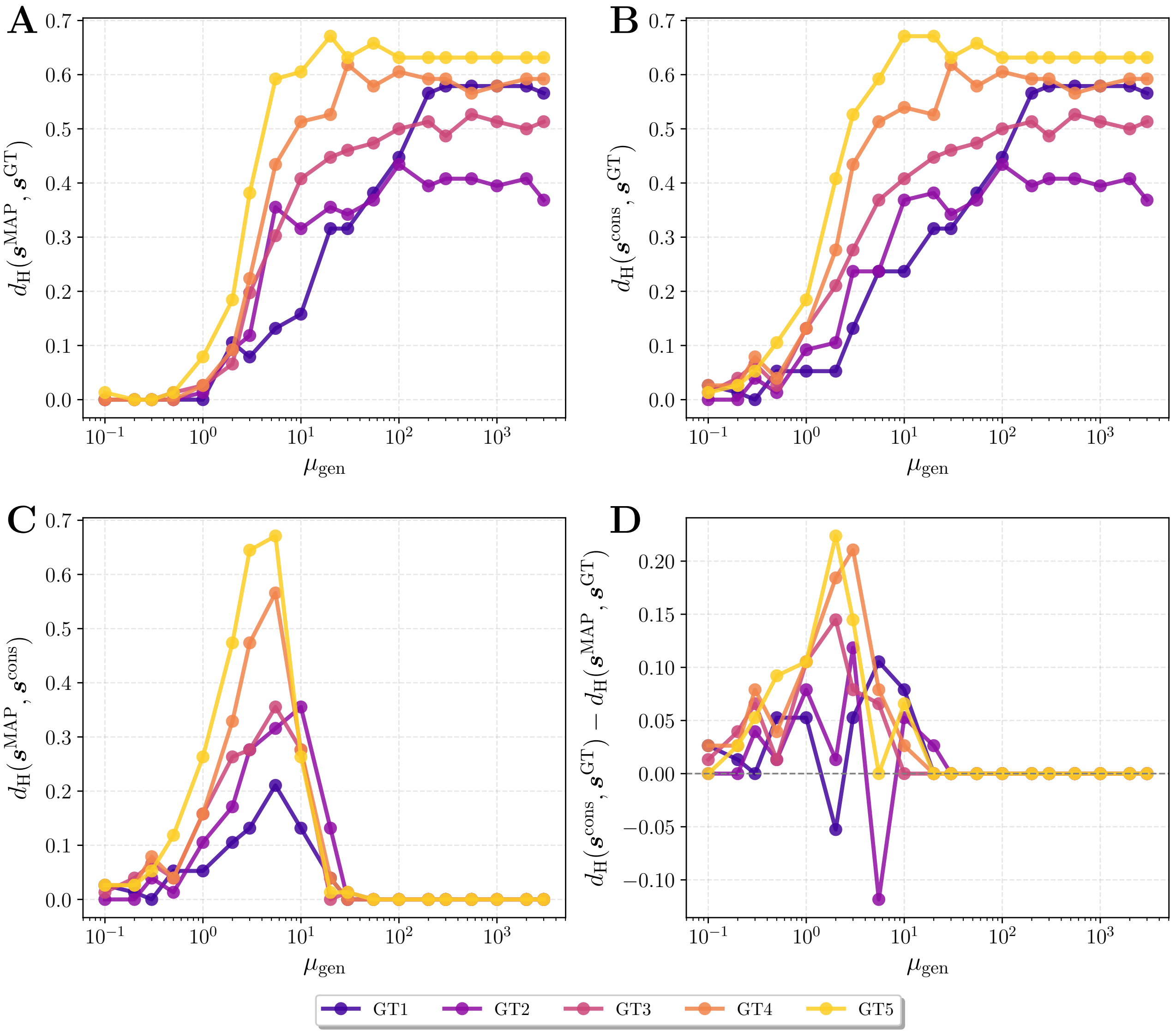


Supplementary Figure 13. **Effect of evolutionary time $\mu_{\mathrm{gen}}$ and root mutability on ASR accuracy. (DBD family)** Normalized Hamming distance plots between Maximum a Posteriori reconstructed ancestor $\boldsymbol{s}^{\mathrm{MAP}}$, GT ancestor $\boldsymbol{s}^{GT}$, and consensus sequence of the leaves $\boldsymbol{s}^{\mathrm{cons}}$ as a function of the number of attempted sweeps $\mu_{\mathrm{gen}}$. (A) Hamming distance between MAP and GT. (B) Hamming distance between consensus sequence from the leaves and GT. (C) Hamming distance between MAP reconstruction and the consensus from the leaves. (D) Difference between the Hamming distances $d_H(\boldsymbol{s}^{\mathrm{cons}}, \boldsymbol{s}^{GT}) - d_H(\boldsymbol{s}^{\mathrm{MAP}}, \boldsymbol{s}^{GT})$. Positive values mean MAP reconstruction is closer to GT than the leaf-consensus.

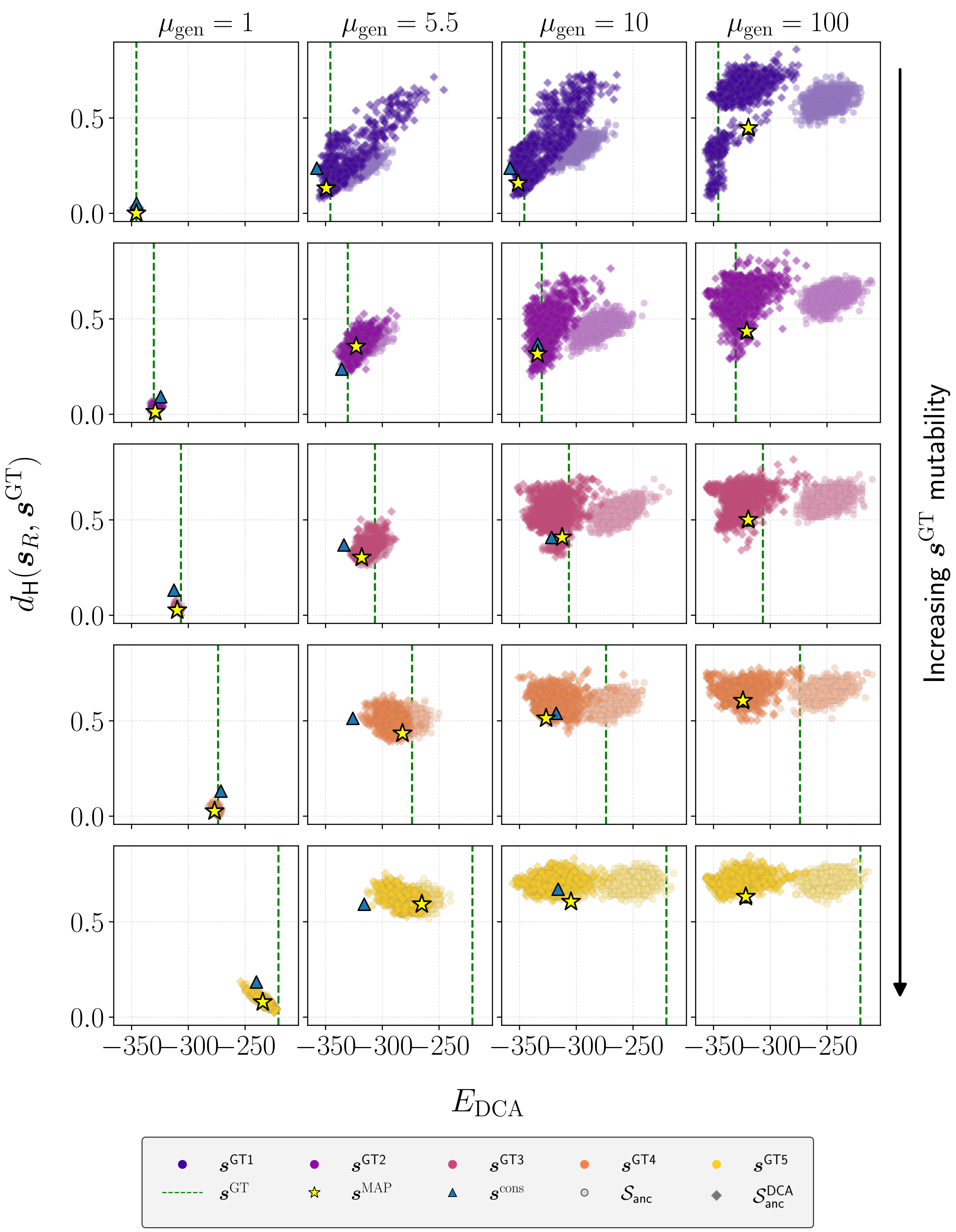


Supplementary Figure 14. **Quantitative assessment of ancestral sequence reconstructions. (DBD family)** Comparison of $\boldsymbol{s}^{GT}$, $\boldsymbol{s}^{\text{MAP}}$, $\boldsymbol{s}^{\text{cons}}$, ancestral samples $\boldsymbol{\mathcal{S}}_{\text{anc}}$ from the site-independent posterior $\mathbb{P}_R$, and 'reshuffled' ancestral samples $\boldsymbol{\mathcal{S}}_{\text{anc}}^{\text{DCA}}$ across evolutionary times and GT roots. Sequences are evaluated using Hamming distance to $\boldsymbol{s}^{GT}$ and DCA energy.

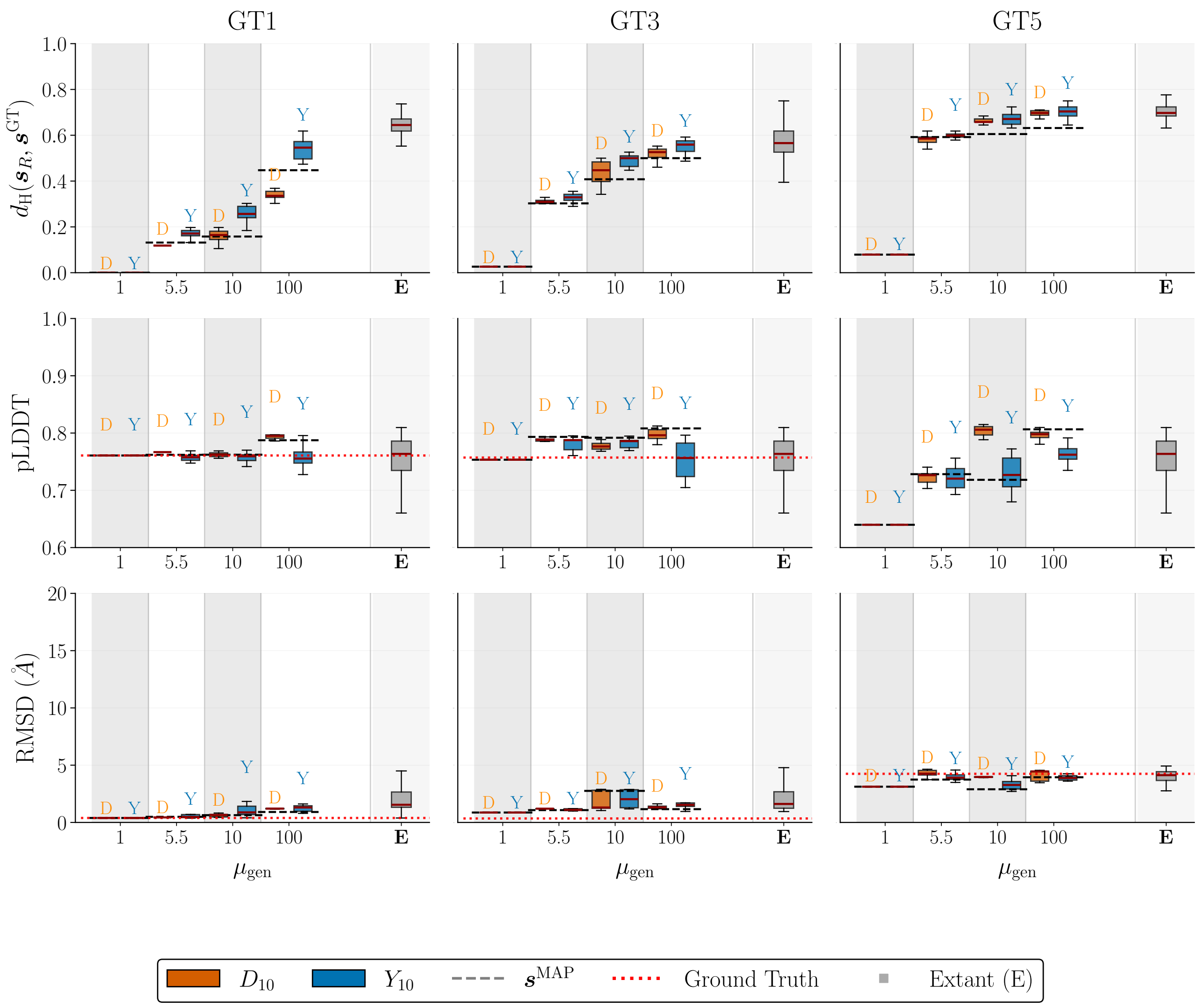


Supplementary Figure 15. **Quantifying proximity of the ancestral samples to $\boldsymbol{s}^{GT}$. (DBD family)** Box-plots of three different reconstruction quality metrics for the extant sequences, $\mathcal{N} = 10$ top-ranking sequences from $\boldsymbol{\mathcal{S}}_{\mathrm{anc}}$ ($\mathrm{Y}_{10}$) and $\boldsymbol{\mathcal{S}}_{\mathrm{anc}}^{\mathrm{DCA}}$ ($\mathrm{D}_{10}$), in terms of site-independent posterior probability $\mathbb{P}_R$. Results are shown for three different roots as a function of evolutionary time $\mu_{\mathrm{gen}}$. First row shows the hamming distance to the $\boldsymbol{s}^{GT}$, with black dotted line representing the hamming distance between $\boldsymbol{s}^{\mathrm{MAP}}$ and $\boldsymbol{s}^{GT}$. Second row is the ESMFold sequence average pLDDT of candidate ancestors, with the red dotted line being the pLDDT of $\boldsymbol{s}^{GT}$. Third row displays the RMSD between ESMFold structural estimates and AlphaFold accurate estimation of the GT structure. Here the red dotted line displays the lower bound of RMSD obtained comparing the stuctural estimates of ESMFold and AlphaFold over $\boldsymbol{s}^{GT}$.

---


[1] EddyRivasLab/hmmer, The Eddy/Rivas Laboratory (2026).
[2] Iqtree/iqtree3, iqtree (2025).
[3] O. Kozlov, Amkozlov/raxml-ng (2026).
[4] S. Guindon, Stephaneguindon/phyml (2026).
[5] L. Nesterenko, L. Blassel, P. Veber, B. Boussau, and L. Jacob, Phyloformer: Fast, Accurate, and Versatile Phylogenetic Reconstruction with Deep Neural Networks, Molecular Biology and Evolution **42**, msaf051 (2025).
[6] M. Price, Morgannprice/fasttree (2025).
[7] S. Whelan and N. Goldman, A general empirical model of protein evolution derived from multiple protein families using a

maximum-likelihood approach, Molecular Biology and Evolution **18**, 691 (2001).

[8] J. Huerta-Cepas, F. Serra, and P. Bork, ETE 3: Reconstruction, Analysis, and Visualization of Phylogenomic Data, Molecular Biology and Evolution **33**, 1635 (2016).

[9] L. Rosset, R. Netti, A. P. Muntoni, M. Weigt, and F. Zamponi, adabmDCA 2.0 – a flexible but easy-to-use package for Direct Coupling Analysis, 2501.18456.
